# SpaFlow depicts the dynamics of ligand-receptor interaction in spatial transcriptomics data

**DOI:** 10.64898/2026.04.17.719264

**Authors:** Hu Chen, Xiaoying Wang, Yuhan Sun, Natalia Del Pilar Vanegas, Jhonny Rodriguez, Ghobashi Ahmed, Anjun Ma, Ana L. Mora, Mauricio Rojas, Qin Ma

## Abstract

Spatial transcriptomics (ST) enables the study of cell-cell communication in native tissue context, but current methods for the ligand-receptor interaction (LRI) inference generally rely on static, distance-based assumptions. Here we present SpaFlow, a reaction-diffusion framework that models ligand diffusion, binding, dissociation, production and degradation to infer spatially resolved LRI activity and hotspots from ST data. Across paired 10x Visium and CosMx metastatic renal cell carcinoma datasets, SpaFlow outperformed existing methods in recovering spatially coherent LRIs, with inferred LRI activity showing stronger association with downstream signaling. In hepatocellular carcinoma after neoadjuvant immunotherapy, SpaFlow identified CXCL12-CXCR4 hotspots enriched at immune-rich tumor boundaries in responders. In aging mouse heart, SpaFlow resolved niche-specific pro-fibrotic and senescence-associated signaling, highlighting Postn-Itgav/Itgb5 as an additional pro-fibrotic axis and Angptl2-Pirb as a candidate mediator of inter-niche senescence-related communication. In human idiopathic pulmonary fibrosis lung, SpaFlow localized CXCL12-CXCR4 signaling between adventitial fibroblasts and CD4 T cells, CD8 T cells, and B cells in the fibrotic surrounding regions. Together, SpaFlow provides a physically informed framework for quantifying spatially constrained cell-cell communication and mechanistically interpreting signaling patterns in complex tissues.

## Main

Cell-cell communication (CCC) is essential for tissue development, homeostasis, repair and disease. Spatial transcriptomics (ST) enables transcriptome profiling in intact tissue and has created new opportunities to study how cellular organization shapes tissue function and pathology^1,2^. A major application of ST is the inference of CCC, much of which is mediated by ligand-receptor interactions (LRIs). In these processes, ligands secreted or presented by one cell engage receptors on nearby cells, forming molecular edges in intercellular signaling networks that regulate immunity, fibrosis, development and tumor progression^3^. Compared with single-cell RNA sequencing (scRNA-seq), which lacks native spatial context, ST provides a more suitable basis for CCC analysis because it constrains candidate interactions by physical proximity and tissue architecture. This is particularly important because scRNA-seq-based CCC analysis can identify ligand-receptor (LR) pairs that are transcriptionally compatible but spatially implausible, thereby increasing false-positive discoveries.

Several computational methods have been developed to infer CCC from ST data. Among them, COMMOT^4^ and SpatialDM^5^ are two widely used and methodologically distinct frameworks for spatially resolved LRI inference. COMMOT models sender-receiver communication using collective optimal transport, inferring directional communication by coupling ligand-producing and receptor-expressing locations under spatial and signaling constraints. SpatialDM identifies interacting LRIs and local communication hotspots by quantifying spatial association between ligand and receptor expression using global and local Moran’s R statistics, thereby detecting spatially enriched interaction patterns across tissue locations. Together, these methods have advanced the analysis of spatial communication by moving beyond cell-type-level summaries and enabling interaction mapping directly in tissue space. However, both rely on a predefined spatial neighborhood or distance-dependent weighting scheme to constrain communication. In COMMOT, signaling is restricted by transport costs over a local spatial domain. In SpatialDM, spatial association is evaluated using a distance-weighted kernel, such as a radial basis function, which likewise requires a user-defined interaction scale. As a result, both methods treat LRIs largely as static, distance-based events.

This assumption can be limiting for diffusible signaling molecules. In tissues, ligand signals are shaped not only by proximity, but also by diffusion, and local consumption. Ligands may spread beyond an assumed interaction radius, accumulate in permissive regions or be depleted in receptor-rich areas that act as local sinks. Shared ligands and receptors can further introduce competition among interacting partners. Consequently, the effective range and intensity of signaling depend on both molecular kinetics and tissue topology, rather than on distance alone. Methods based primarily on spatial correlation or fixed distance cutoffs may therefore miss biologically meaningful signaling patterns, particularly when hotspots are shaped by the coupled effects of diffusion, reaction kinetics and tissue topology.

Reaction-diffusion models provide a natural framework for describing these processes. In principle, they can jointly represent ligand spreading, receptor engagement, complex formation and turnover within a unified mechanistic model. This offers an attractive alternative to static scoring approaches because it links observed spatial patterns to explicit physical processes. In practice, however, applying reaction-diffusion models to ST data is challenging. Analytical solutions are rarely available in realistic tissues with irregular geometry and heterogeneous cell spacing, and direct numerical simulation can be computationally expensive when applied across many spatial locations and LR pairs. These challenges are further amplified by the diversity of ST platforms, which range from spot-based assays such as 10x Visium to single-cell and subcellular imaging-based methods, and therefore require a representation that is flexible across different spatial resolutions and sampling schemes.

Here we present SpaFlow, a graph-based framework for CCC inference that models LRIs as dynamic processes over the spatial organization of spots or cells, which we collectively refer to as spots. In SpaFlow, each spatial unit is represented as a node in a graph, and neighboring units are connected by edges, enabling tissue architecture to be captured without imposing a regular grid. Ligand diffusion is represented as flux across graph edges using a discrete form of Fick’s first law, allowing molecules to move from regions of high concentration to regions of low concentration. LR binding and dissociation follows mass-action kinetics, linking local ligand availability and receptor abundance to the formation of signaling complexes. Production and degradation terms account for secretion and molecular clearance, and effective ligand and receptor concentrations are used to capture competition among shared interaction partners. Together, these components define a system of ordinary differential equations (ODEs) that describes diffusion, reaction, production and degradation on the spatial graph.

This formulation replaces an imposed interaction radius with a mechanistically grounded description of how signaling propagates through tissue and enables both interaction scoring and hotspot detection within a unified framework. Because SpaFlow explicitly models diffusion, binding, dissociation, production and degradation in the coupled ligand-receptor-complex system, it also supports mechanistic interrogation of the processes that shape spatial signaling patterns. Across six paired 10x Visium and CosMx metastatic renal cell carcinoma (mRCC) datasets^6^, SpaFlow outperformed existing methods in recovering spatially coherent and regulatory-relevant LRIs. Application to hepatocellular carcinoma (HCC), aging mouse heart and human idiopathic pulmonary fibrosis (IPF) lung datasets further identified biologically meaningful hotspots and condition-specific signaling programs associated with tumor-immune boundaries, cardiac aging and fibrotic microenvironments. By treating CCC as a dynamic biochemical process rather than a static distance-based score, SpaFlow provides a physically informed framework for studying how signaling emerges within native tissue architecture.

## Results

### Overview of SpaFlow

SpaFlow is designed to infer spatially resolved LRI activity from ST data by explicitly modeling how signaling is shaped by tissue architecture (**Fig. 1**). SpaFlow takes ST from platforms such as 10x Visium and CosMx together with a curated LR database CellChatDB^7^ as input. Because SpaFlow models LRIs as a dynamical process in tissue, ligand and receptor concentrations for each LR pair are initialized from the expression levels of their corresponding genes, following a common assumption in transcriptomics-based studies of cell-cell communication (CCC), whereas the concentration of their complex, is initialized to zero because it is not directly observed and cannot be reliably approximated from the input data. Spots are represented as nodes in a spatial graph constructed using a k-nearest-neighbor strategy, which provides a tractable approximation for signaling dynamics in irregularly sampled tissues. Because SpaFlow explicitly models ligand diffusion across tissue space, the current framework is primarily designed for secreted signaling, in which diffusible ligands mediate communication across neighboring or more distant cells.

**Fig. 1.**
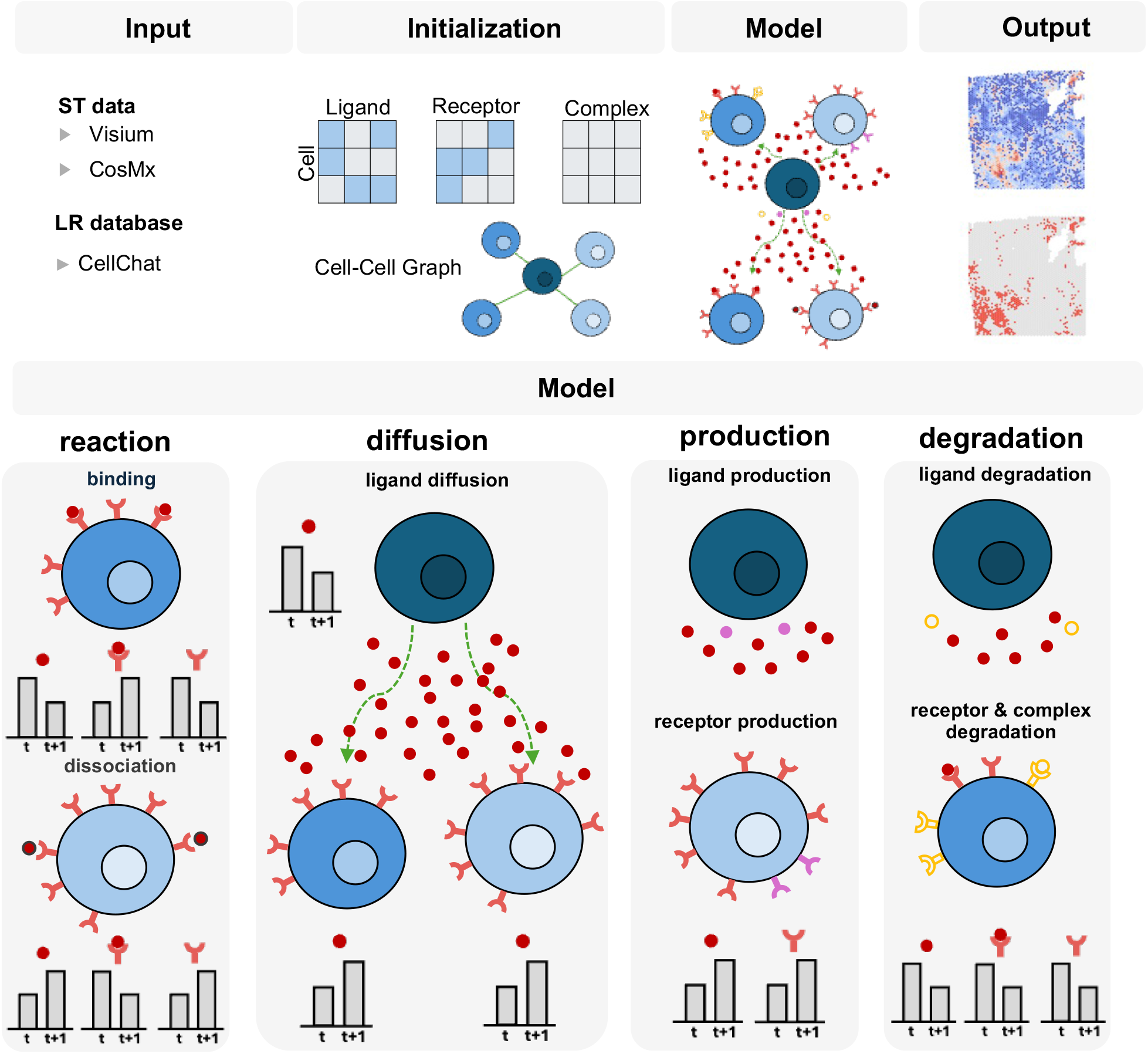
Overview of SpaFlow. SpaFlow takes ST data together with a prior LR database as input. Ligand and receptor concentrations are initialized from gene expression, whereas LR complex concentration is initialized to zero. Cells or spots are represented as nodes in a spatial neighbor graph, and each LRI is modeled as a graph-based dynamical system incorporating five coupled processes: binding, dissociation, diffusion, production and degradation. The system is iteratively updated until it reaches a steady state. Outputs include spatial maps of complex concentration, which quantify interaction activity, and signaling hotspots identified by spatial permutation testing.

Using this graph representation, SpaFlow models each LRI as a spatially constrained dynamical system and iteratively updates ligand, receptor and complex concentrations until the system reaches a steady state. This formulation enables the model to account for both local molecular availability and neighborhood structure, thereby capturing communication patterns that depend on spatial context rather than expression co-occurrence alone. SpaFlow therefore provides a quantitative framework for estimating where specific LRIs are active within tissue.

At convergence, SpaFlow outputs spatial maps of complex concentration, which serve as quantitative estimates of LRI activity across tissue locations. These interaction scores can be analyzed at the level of individual LRIs or aggregated into pathway-level activities for downstream comparisons across samples or clusters. In addition, SpaFlow identifies hotspots through spatial permutation testing, enabling detection of significantly localized signaling regions within tissue. Together, these features allow SpaFlow to quantify both the magnitude and the spatial localization of LRI activity in intact tissues.

### Mathematical formulation, benchmarking and ablation analysis of SpaFlow

We next made this framework explicit as a system of ODEs describing the update of ligand, receptor and complex concentrations for each LRI on the spatial graph (**Fig. 2a**). In these equations, ligand concentration is governed by diffusion between neighboring nodes together with local production, receptor binding, complex dissociation and degradation; receptor concentration is governed by production, ligand binding, complex dissociation and degradation; and complex concentration is governed by LR binding, dissociation and degradation. SpaFlow further defines effective ligand and receptor concentrations to account for competition among ligands and receptors participating in multiple interactions, thereby allowing each LRI to be evaluated in the context of shared molecular usage.

**Fig. 2.**
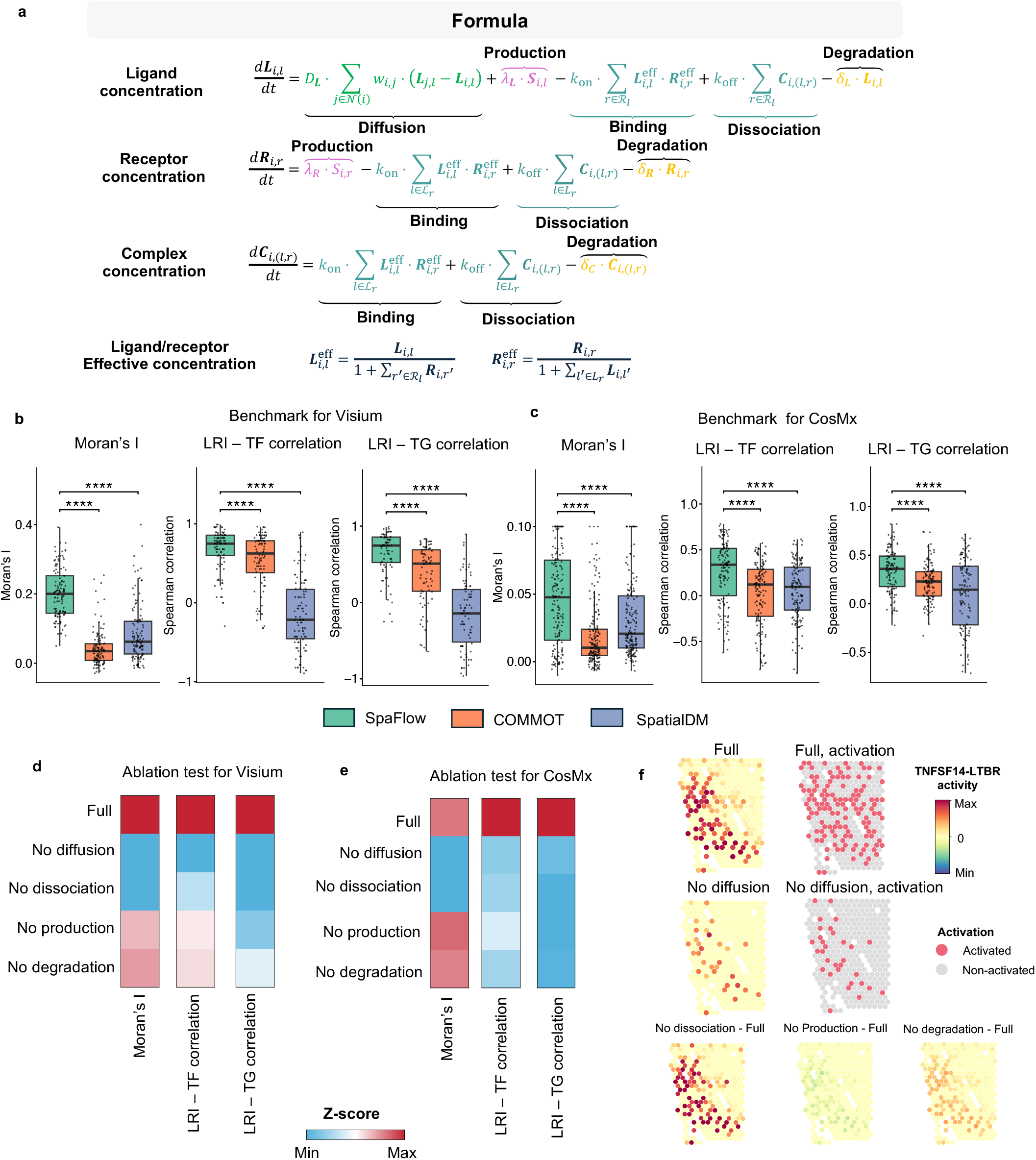
ODE formulation, performance evaluation and ablation analysis of SpaFlow. **a**, System of ODEs defining the dynamics of ligand, receptor and complex concentrations for each LRI on the spatial graph. Effective ligand and receptor concentrations are used to account for competition among shared interaction partners. **b**, Benchmarking on a representative Visium sample, comparing SpaFlow, COMMOT and SpatialDM using Moran’s I and Spearman’s correlations between inferred LRI activity and TF or TG activity. **c**, Benchmarking on a representative CosMx sample using the same evaluation metrics as in **b**. **d**, Ablation analysis on a representative Visium sample showing the effect of removing diffusion, dissociation, production or degradation from the full model. Heatmap values represent the mean metric across all LRIs in the sample, and color indicates Z-scored performance within each metric. **e**, Ablation analysis on a representative CosMx sample using the same evaluation metrics as in **d**. **f**, Spatial maps of TNFSF14-LTBR activity in a highlighted region from one representative Visium sample, shown for the full model and the no diffusion variant. For the no dissociation, no production and no degradation variants, only difference maps relative to the full model are shown to highlight local changes.

We then benchmarked SpaFlow against COMMOT and SpatialDM using paired Visium and CosMx datasets from the mRCC study, comprising six Visium samples and six CosMx samples in total. Performance was evaluated using three complementary criteria: spatial autocorrelation of inferred LRI activity, measured by Moran’s I, and concordance with downstream singaling, assessed by Spearman’s correlations between LRI activity and transcription factor (TF) activity or target gene (TG) activity (**Fig. 2b, c**). In the representative Visium sample, SpaFlow achieved higher Moran’s I than both comparison methods, indicating improved recovery of spatially coherent communication signals. SpaFlow also showed stronger correlations with both TF and TG activity, suggesting that its inferred LRI scores more closely reflect downstream regulatory effects. Similar results were observed in another representative CosMx sample, where SpaFlow consistently outperformed COMMOT and SpatialDM across all three metrics. Consistent with these representative examples, benchmarking across all six Visium samples and all six CosMx samples showed the same overall performance advantage for SpaFlow (**Supplementary Fig. 1, Supplementary Tabel. 1**). Together, these results indicate that SpaFlow more effectively captures both the spatial organization and the regulatory consequences of cell-cell communication across distinct ST platforms.

To determine the contribution of individual model components, we performed ablation analysis by systematically removing diffusion, dissociation, production or degradation from the full model. In representative Visium and CosMx samples, the full model showed the strongest overall performance across Moran’s I and the two regulatory concordance metrics (**Fig. 2d, e**). Removing diffusion or dissociation generally led to the largest reductions in performance, whereas removing production or degradation had comparatively smaller effects and in several samples yielded results close to those of the full model. This pattern was most evident for Moran’s I and was also observed for correlations with TF and TG activity, for which the no-diffusion and no-dissociation variants generally showed larger performance losses than the no-production and no-degradation variants. The same overall pattern was consistently observed across all six Visium samples and all six CosMx samples from the mRCC study (**Supplementary Fig. 2, Supplementary Tabel. 2**). Together, these results support the full SpaFlow formulation and indicate that diffusion and dissociation make the strongest contributions to model performance under these evaluation settings, whereas production and degradation make comparatively smaller contributions.

To illustrate the effects of removing individual terms, we examined TNFSF14-LTBR in a representative Visium sample (**Fig. 2f**). Removing diffusion reduced inferred LRI activity in spots surrounding regions with high interaction activity, consistent with the role of diffusion in propagating ligand signals across neighboring locations. Removing dissociation markedly increased inferred LRI activity, consistent with reduced complex dissociation and accumulation of bound complexes. By contrast, removing production or degradation shifted LRI activity downward or upward, respectively, reflecting the role of these processes in maintaining dynamic balance within the system.

### SpaFlow identifies CXCL12-CXCR4 hotspots at immune-enriched tumor boundaries in responder HCC following immunotherapy

Using SpaFlow, we systematically compared LRIs across six advanced HCC surgical resection samples profiled by 10x Visium following neoadjuvant immunotherapy^8^, including three responders and three non-responders, to define conserved spatial communication programs associated with therapeutic response. Only one LR pair, CXCL12-CXCR4, was shared across responder samples, whereas most detected LRIs were sample-specific, highlighting both conserved and patient-specific spatial signaling features within the HCC microenvironment (**Fig. 3a**). Given its consistency and abundance in responders, we next focused on the spatial organization and functional relevance of CXCL12-CXCR4 hotspots.

**Fig. 3.**
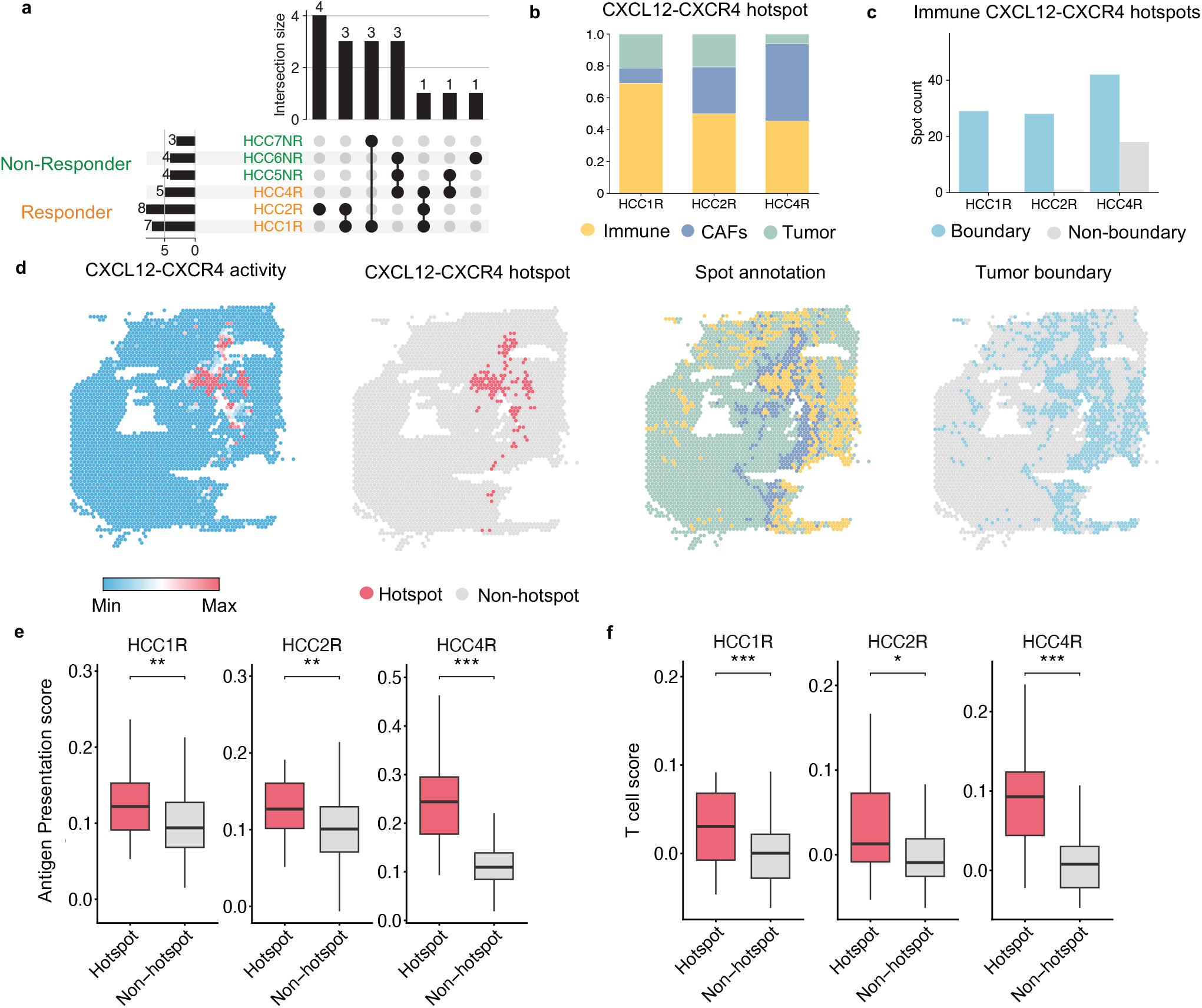
SpaFlow identifies CXCL12-CXCR4 hotspots at immune-enriched tumor boundaries in responder HCC following neoadjuvant immunotherapy. **a**, UpSet plot showing the overlap of SpaFlow-detected LRIs across six advanced HCC samples profiled by 10x Visium after neoadjuvant cabozantinib plus nivolumab treatment. **b**, Spot annotation of CXCL12-CXCR4 hotspots in the three responder samples. **c**, Numbers of immune CXCL12-CXCR4 hotspots located at tumor boundaries or non-boundary regions. **d**, Spatial maps of a representative sample HCC4R showing CXCL12-CXCR4 activity, CXCL12-CXCR4 hotspot distribution, spot annotation and tumor-boundary regions. **e**, Antigen-presentation scores in CXCL12-CXCR4 hotspot versus non-hotspot spots. **f**, T cell scores in CXCL12-CXCR4 hotspot versus non-hotspot spots.

Compositional analysis of spot annotations showed that CXCL12-CXCR4 hotspots were predominantly associated with immune spots across all responder samples, which was further supported by spatial mapping of hotspot regions onto the tissue sections (**Fig. 3b, e, Supplementary Tabel. 3**). In parallel, quantification of immune CXCL12-CXCR4 hotspots demonstrated that these regions were consistently enriched at tumor boundaries across all three responder samples, again in agreement with their spatial localization patterns (**Fig. 3c, e, Supplementary Tabel. 4**). Together, these findings indicate that CXCL12-CXCR4 hotspots are not randomly distributed, but instead preferentially localize to immune-enriched boundary niches in responder tumors.

To assess the functional relevance of these SpaFlow-identified CXCL12-CXCR4 hotspots, we compared pathway activities between hotspot and non-hotspot spots. CXCL12-CXCR4 hotspots exhibited significantly higher antigen presentation scores across all three responder samples (**Fig. 3e, Supplementary Tabel. 5**). T cell scores were likewise significantly elevated in hotspot regions relative to non-hotspot regions (**Fig. 3f, Supplementary Tabel. 6**). Collectively, these findings suggest that in responders, the CXCL12-CXCR4 axis is associated with spatially organized tumor-margin niches characterized by immune cell enrichment, enhanced antigen presentation, and increased T cell activity. This spatial-functional coupling may represent an important microenvironmental feature that supports effective anti-tumor immune responses following neoadjuvant therapy.

### SpaFlow reveals pro-fibrotic signaling and SASP-associated inter-niche communication in the aging mouse heart

We applied SpaFlow to ten 10x Visium mouse heart samples, including five young mice (3 months) and five old mice (18 months), to identify aging-associated changes in intercellular communication in the mouse heart. To simplify differential analysis, we first aggregated LRIs into pathway-level activities and hotspots and then compared pathway activity between young and old hearts at the sample level. This analysis identified eight secreted signaling pathways that were significantly altered with aging (**Fig. 4a, Supplementary Tabel. 7**). We next examined these changes at niche resolution. COMPLEMENT activity was significantly enriched in niches 0, 1 and 7, whereas PERIOSTIN activity was significantly enriched in niches 2 and 7 (**Fig. 4b, Supplementary Tabel. 8**). Notably, niche 7 was previously defined in the original study as a pro-inflammatory and pro-fibrotic vascular niche in the aging heart^9^.

**Fig. 4.**
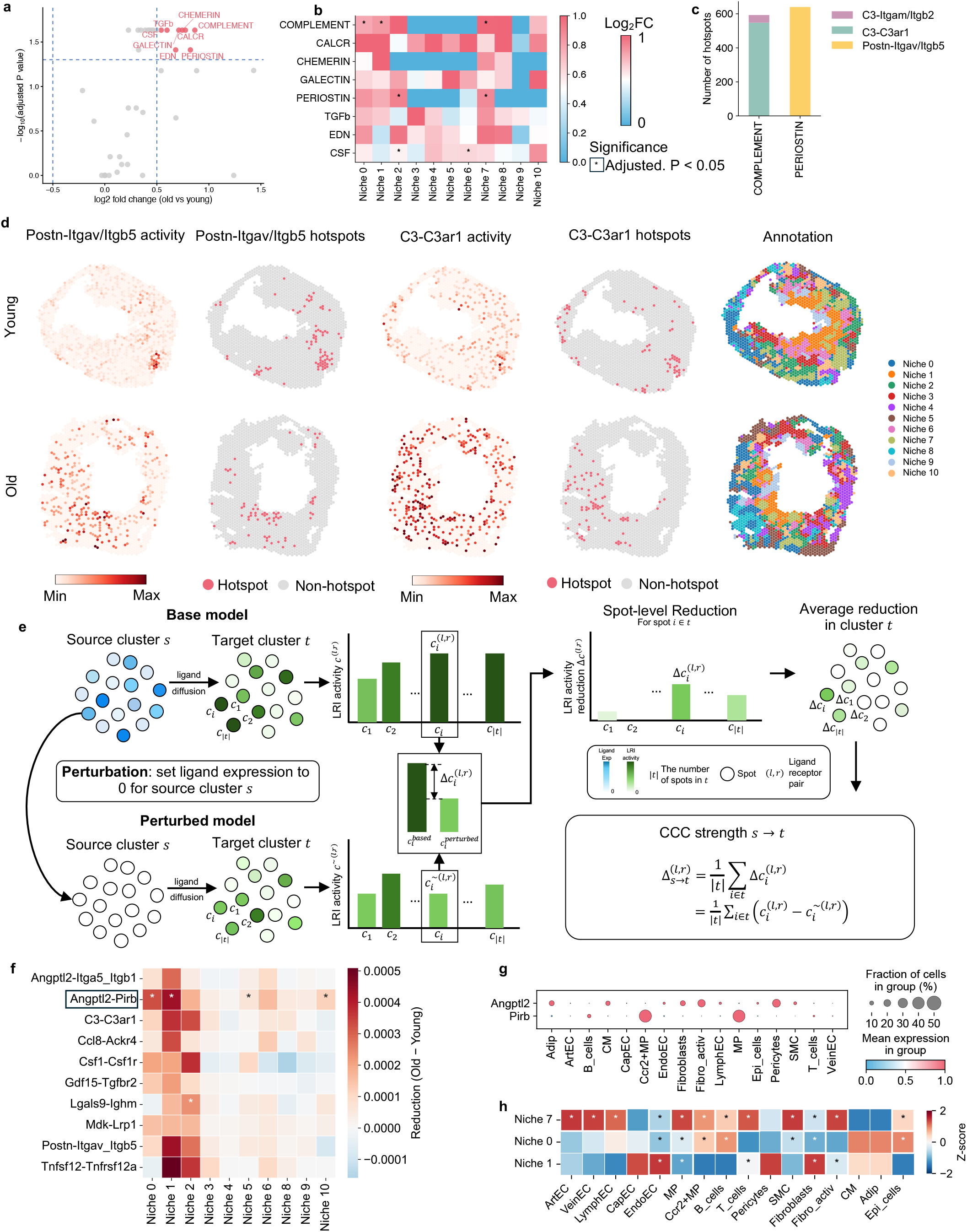
SpaFlow identified aging-related pro-fibrotic and senescence-related LRIs in mouse heart. **a**, Differential pathway activity analysis between old and young mouse hearts at the sample level. Significant pathways are defined as those with |log2FC| > 0.5 and adjusted P < 0.05, assessed using a two-sided Wilcoxon rank-sum test with Benjamini-Hochberg correction. **b**, Log2FC of pathway activity between old and young mouse hearts at the niche level. Significance is indicated by * (adjusted P < 0.05), assessed using a two-sided Wilcoxon rank-sum test with Benjamini-Hochberg correction across niches. **c**, Bar plots showing the contribution of hotspot-associated LRIs to selected pathways. **d**, Spatial maps of Postn-Itgav/Itgb5 activity, C3-C3ar1 activity, senescence score and spot annotations in one representative young sample and one representative old sample. **e**, Schematic illustration of the in silico ligand perturbation analysis. **f**, Perturbation analysis of niche 7. Color indicates the change in interaction activity (old - young), with lower values indicating greater ligand influence in old samples. The top 10 LRIs are ranked by average change across perturbed samples. Significance is assessed using a two-sided Wilcoxon rank-sum test with Benjamini-Hochberg correction; adjusted P < 0.05. **g**, scRNA-seq gene expression dot plot for Angplt2 and Pirb. **h**, Cell-type proportion heatmap showing enrichment of cell types across niches. Data are Z-score transformed across niches for each cell type. Significance is assessed using Fisher’s exact test with Benjamini-Hochberg correction; adjusted P < 0.05.

To identify the LRIs underlying these pathway signals, we decomposed pathway hotspots into individual LR contributions. The dominant interactions were C3-C3ar1 for COMPLEMENT and Postn-Itgav/Itgb5^10^ for PERIOSTIN (**Fig. 4c, Supplementary Tabel. 9**). Representative spatial maps from young and old hearts showed that both pathway activity and hotspot signals were most prominent in niche 7 (**Fig. 4d**). C3-C3ar1 was identified in the original study as a pro-fibrotic signal, likely reflecting fibroblast-macrophage crosstalk. We further suggested that Postn-Itgav/Itgb5 represents an additional pro-fibrotic signaling axis, consistent with its established role in fibrosis.

Niche 7 also showed the highest senescence burden among all cardiac niches in the original study, supporting its role as a major SASP-associated niche in the aged heart. To identify LRIs potentially involved in the spread of senescence-associated signals, we performed in silico ligand perturbation analysis to quantify the influence of niche 7 as a source niche on other niches (**Fig. 4e**). Specifically, for each LR pair (*l, r*), we defined the CCC strength 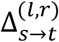 from source cluster *s* to target cluster *t* as the average reduction in LRI activity between the original and perturbed conditions across spots in the target cluster. Comparing this reduction between old and young hearts identified Angptl2-Pirb^11^ as an LR pair with a stronger perturbation-induced decrease in LRI activity in niches 0 and 1 in old hearts than in young hearts (**Fig. 4f, Supplementary Tabel. 10**), suggesting that this signaling axis may contribute to the propagation of senescence-associated signals from niche 7, where fibroblasts in niche 7 secrete Angptl2 and macrophages in niches 0 and 1 receive it through Pirb, potentially promoting macrophage senescence^12,13^.

Consistent with this result, matched scRNA-seq data from the same study showed that fibroblasts and activated fibroblasts were the principal sources of Angptl2, whereas macrophages and Ccr2+ macrophages were the predominant receiver populations expressed receptor Pirb (**Fig. 4g**). Cell-type proportion heatmaps further showed that niche 0 was enriched for macrophages and Ccr2+ macrophages, niche 1 for macrophages, and niche 7 for fibroblasts and activated fibroblasts (**Fig. 4h, Supplementary Tabel. 11**). Together, these results identify a pro-fibrotic and SASP-related signaling program centered on niche 7 that may influence adjacent niches during cardiac aging through fibroblasts and macrophages communications.

### SpaFlow localized CXCL12-CXCR4 signaling in the fibrotic surrounding region of human IPF lungs

We applied SpaFlow to six 10x Visium spatial transcriptomic samples from human lung tissue, including two healthy controls and paired IPF upper lobe (UL) and IPF lower lobe (LL) samples from two patients. CXCL12-CXCR4 showed the highest number of hotspots in IPF UL plus IPF LL samples and the second highest proportion of hotspots for IPF UL plus IPF LL samples relative to hotspots in all samples (**Fig. 5b, Supplementary Tabel. 12**). Spatial plots for spot annotation, CXCL12-CXCR4 activity, and corresponding hotspots are shown (**Fig. 5a**), where lung fibrosis spots occupy the largest fraction of hotspots for IPF samples compared with healthy controls (**Fig. 5c, Supplementary Tabel. 13**). CXCL12-CXCR4 activity on lung fibrosis hotspots increases from control to IPF UL and IPF LL, consistent with previous biological findings^14-16^, whereas COMMOT and SpatialDM cannot detect this difference (**Fig. 5d, Supplementary Tabel. 14**).

**Fig. 5.**
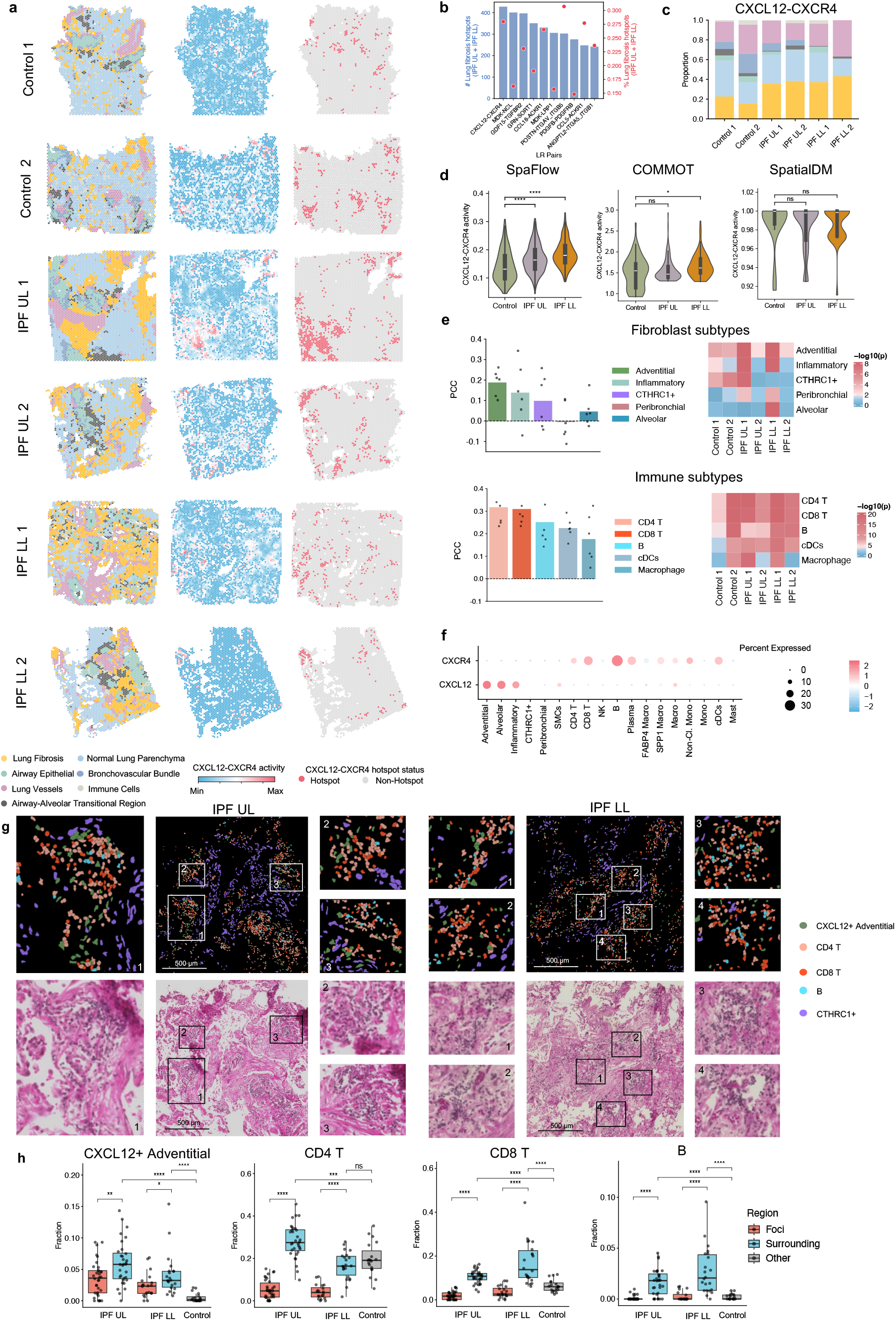
SpaFlow identifies CXCL12-CXCR4 as a key LRI in fibrotic surrounding region of human IPF lungs. **a**, Spatial maps for six 10x Visium samples (two control, two IPF UL and two IPF LL). Left, spot annotations. Middle, CXCL12-CXCR4 activity. Right, hotspot status for CXCL12-CXCR4 activity. **b**, Number of CXCL12-CXCR4 hotspots in IPF UL and IPF LL samples, and the proportion of hotspots in IPF samples (UL + LL) relative to all samples (control + IPF UL + IPF LL). **c**, Distribution of spot annotations within hotspots for each sample. **d**, CXCL12-CXCR4 activity in hotspots across conditions (control, IPF UL and IPF LL) estimated by SpaFlow, COMMOT and SpatialDM. Significance is assessed using a two-sided Wilcoxon rank-sum test; ns, P > 0.05;, P ≤ 0.05; **, P ≤ 0.01; ***, P ≤ 0.001; ****, P ≤ 0.0001. **e**, Left, Pearson correlation between CXCL12-CXCR4 activity and cell-type abundance across fibrotic spots; the top five fibroblast subtypes and top five immune subtypes are shown. Right, heatmap showing the significance of the selected subtypes in each sample. **f**, scRNA-seq gene expression dot plot of CXCL12 and CXCR4 across fibroblast and immune-cell subtypes. **g**, Representative Xenium images from IPF UL (left) and IPF LL (right), showing colocalization of CXCL12+ adventitial fibroblasts with CD4 T cells, CD8 T cells and B cells; corresponding H&E images are shown below. **h**, Quantification comparing fibrotic foci with fibrotic surrounding regions in IPF UL and IPF LL, and fibrotic surrounding regions with normal control regions. Dots indicate pathologist-annotated regions. Significance is assessed using a two-sided Wilcoxon rank-sum test; ns, P > 0.05; *, P ≤ 0.05; **, P ≤ 0.01; ***, P ≤ 0.001; ****, P ≤ 0.0001.

Next, we asked which fibroblast and immune subtypes contribute to CXCL12-CXCR4 signaling in lung fibrosis spots. Prior scRNA-seq studies support fibroblast immune crosstalk mediated by CXCL12-CXCR4, including T cells, B cells^14^, and plasma cells^17^, but they do not resolve fibroblast subtypes and lack spatial context. We therefore used paired scRNA seq data to deconvolve Visium spots with cell2location^18^ and computed Pearson correlations between CXCL12-CXCR4 activity and inferred cell type abundance **(Fig. 5e, Supplementary Tabel. 15)**. Among fibroblast subtypes, adventitial fibroblasts ranked first, showing the highest average correlation and significance across all six samples. Among immune subtypes, CD4 T cells, CD8 T cells, and B cells were prioritized based on consistently higher correlations and significance across all six samples. Paired scRNA-seq gene expression data further showed that adventitial fibroblasts are the predominant source of CXCL12, whereas B cells, CD4 T cells, and CD8 T cells constitute the major CXCR4-expressing populations (**Fig. 5f**). Together, these results suggest that adventitial fibroblasts engage CD4 T cells, CD8 T cells, and B cells through the CXCL12-CXCR4 axis within lung fibrotic regions.

However, because 10x Visium measures transcript abundance at the spot level, the annotated lung fibrosis region may include both established fibrotic foci and the surrounding tissue. To obtain fine grained spatial validation, we next analyzed Xenium in situ datasets from one healthy control, one IPF UL, and one IPF LL lung sample. In each dataset, we annotated spatial domains corresponding to the fibrotic foci and the surrounding region, whereas in the healthy control we annotated the comparable area as others. The fibrotic foci were dominated by CTHRC1+ fibroblasts, whereas CXCL12+ adventitial fibroblasts were enriched in the perilesional surrounding compartment. Within this region, CXCL12+ adventitial fibroblasts were spatially colocalized with CD4 T cells, CD8 T cells, and B cells, with CD4 T and CD8 T cells being the most abundant lymphocyte populations (**Fig. 5g**). Quantification further showed increased fractions of CXCL12+ adventitial fibroblasts, CD4 T cells, CD8 T cells, and B cells in the fibrotic surrounding region compared with the fibrotic foci (**Fig. 5h, Supplementary Tabel. 16**), supporting a hypothsis in which CXCL12 producing adventitial fibroblasts engage CXCR4 expressing T and B cells at the margins of fibrotic lesions, thereby promoting immune cell recruitment and local inflammatory signaling, this also support the structure of tertiary lymphoid structures exists in IPF samples^19^ because of the accumulation of immune cells.

## Discussion

ST has expanded the study of CCC in intact tissues, but most existing methods still infer LRIs primarily from spatial proximity or expression covariance. Although useful in most cases, such formulations do not explicitly represent the physical processes that shape signaling in tissue, particularly for diffusible ligands. SpaFlow addresses this gap by modeling LRI activity as a reaction-diffusion process on a spatial graph, integrating ligand, diffusion, binding, dissociation, production and degradation within a unified framework. Across paired Visium and CosMx datasets, this formulation improved recovery of spatially coherent LRIs and strengthened concordance with downstream singnaling, indicating that explicit modeling of signaling dynamics can improve CCC inference from ST data.

A central advantage of SpaFlow is that it replaces a predefined interaction radius with a mechanistic description of signal propagation and local consumption. In tissues, the effective range and intensity of signaling are shaped not only by distance, but also by diffusion, molecular turnover, receptor-mediated depletion and competition among shared ligands or receptors. These coupled processes can generate signaling patterns that are not readily captured by static neighborhood definitions or spatial association statistics. Consistent with this interpretation, ablation analysis showed that diffusion and dissociation contributed most strongly to performance, supporting the view that spatial transport and reversible complex formation are major determinants of the inferred signaling landscape.

The biological applications further illustrate the utility of this framework across distinct tissue contexts. In HCC patients after neoadjuvant immunotherapy, SpaFlow identified CXCL12-CXCR4 hotspots at immune-enriched tumor boundaries in responders, linking this axis to localized antigen presentation and T cell activity. In the aging mouse heart, SpaFlow resolved niche-specific pro-fibrotic and inflammatory signaling, highlighted Postn-Itgav/Itgb5 as an additional candidate pro-fibrotic axis and implicated Angptl2-Pirb in SASP-associated inter-niche communication. In IPF, SpaFlow localized CXCL12-CXCR4 signaling to the fibrotic surrounding region, and integration with deconvolution and Xenium data supported a model in which CXCL12-producing adventitial fibroblasts engage CXCR4-expressing lymphocytes at lesion margins. Together, these results show that SpaFlow can define not only which LRIs are active, but also where they are spatially organized and which tissue compartments may underlie disease-associated signaling.

This study has several limitations. SpaFlow is currently designed for secreted signaling and does not explicitly model contact-dependent communication, extracellular matrix sequestration or anisotropic transport barriers. As with most transcriptomics-based CCC methods, transcript abundance is used as a proxy for molecular concentration and activity, which does not capture post-transcriptional regulation, secretion efficiency, receptor localization or protein turnover. In addition, the current implementation uses a fixed graph representation and globally specified kinetic parameters to obtain steady-state solutions, trading biochemical specificity for general applicability and computational tractability. Future extensions could incorporate spatial proteomic measurements, interaction-specific parameterization, matrix-aware transport and time-resolved or perturbational spatial data. More broadly, SpaFlow provides a physically informed framework for moving ST-based CCC analysis beyond descriptive association toward mechanistic inference and should be broadly useful for dissecting signaling programs across spatial omics platforms and disease settings.

## Methods

### Modeling ligand-receptor interaction through ordinary differential equations

SpaFlow takes a gene expression matrix ***X*** ∈ ℝ^*n*×*m*^ and spot coordinates ***S*** ∈ ℝ^*n*×2^ from ST data as input, together with a curated LR database from CellChatDB with signaling type = ‘Secreted Signaling’. Here, *n* denotes the number of spots, *m* denotes the number of genes. To model the dynamic process of LRI, for each LR pair (*l, r*) in spot *i*, we initialize ligand concentration ***L***_*i,l*_ and receptor concentration ***R***_*i,r*_ using their corresponding gene expression, and initialize the complex concentration ***C***_*i*,(*l,r*)_ to zero. Because analytical solutions are typically intractable in realistic tissues, we construct a spot graph ***G*** from the spatial coordinates using *k* nearest neighbors with *k* = 6 in default. Edge weights *w*_*i,j*_ are derived from the symmetrically normalized graph Laplacian:

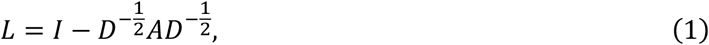

where *A* ∈ ℝ^*n*×*n*^ is the adjacency matrix and *D* ∈ ℝ^*n*×*n*^ is the diagonal degree matrix. *A*_*ij*_ = 1 if spot *i* and *j* are adjacent and *A*_*ij*_ = 0 othewise. Accordingly, the normalized edge weight between adjacent spot *i* and *j* is defined as:

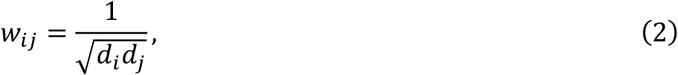

Where *d*_*i*_ and *d*_*j*_ denote the degree of spot *i* and *j*, respectively. This formulation allows diffusion to be represented as flux along graph edges.

Given this discrete formulation, we use the following ODEs to describe the update of ligand, receptor and complex concentrations at each iteration *t*:

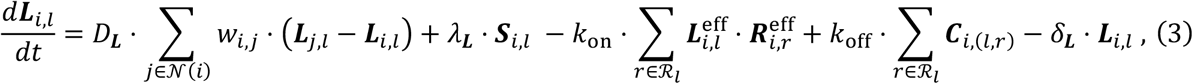

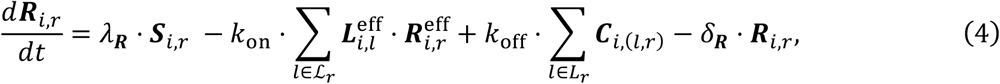

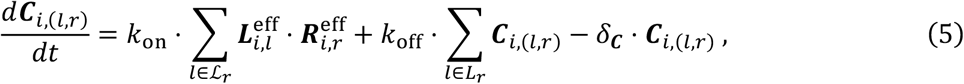

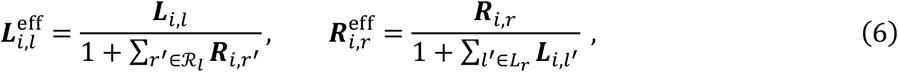

where *D*_***L***_ is the ligand diffusion rate, *k*_*on*_ and *k*_*off*_ are binding and dissociation rates, *λ*_***L***_ and *λ*_***R***_ are the production rates of ligand and receptor. *δ*_***L***_, *δ*_***R***_ and *δ*_***C***_ are the degradation rates of ligand, receptor and complex. 𝒩(*i*) denotes the set of neighbors of spots *i*, meaning that at each time step the ligand diffuses to nearby spots. ℛ_*l*_ denotes all receptors bind to ligand *l*, and ℒ_*r*_ denotes all ligands bind to receptor *r*. Within a spot, multiple receptors can bind the same ligand, and multiple ligands can bind the same receptor, so we use Hill-type competition functions to model the competition. 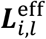 is the effective concentration of ligand *l* after accounting for competing receptors ℛ_*l*_, and 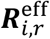 is the effective concentration of receptor *r* after accounting for competing ligands ℒ_*r*_.

We numerically solve this ODE system using a forward Euler scheme with step size Δ*t* . Specifically, at iteration *t*, ligand, receptor and complex concentrations are updated as:

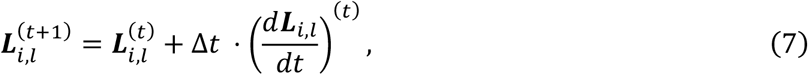

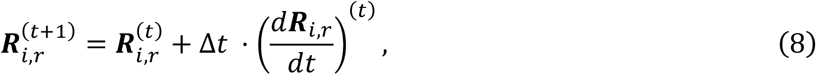

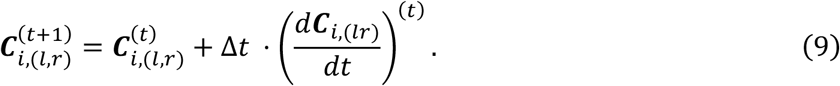

where the right-hand-side derivatives are evaluated using Equation. (3)-(7) at iteration *t*. Here, superscript (*t*) denotes the iteration index of the numerical scheme. We initialize ligand and receptor concentrations from gene expression, initialize complex concentration to zero, and iterate the system until convergence. In practice, we run the simulation for *T* = 500 iterations with Δ*t* = 0.01, and use the final steady-state complex concentration to quantify LRI activity at each spot. All hyperparameters are set to ensure the system reaches a steady state.

### Defining ligand-receptor hotspots via spatial permutation testing

To define hotpots for each LRI, we used a spatial permutation test that breaks spatial structure while keeping gene expression fixed. We first computed the steady state complex concentration *c*_*i*_ at each spot *i* using the original coordinates and the corresponding graph Laplacian. We then generated a null distribution by permuting spot coordinates across spots *B* times, recomputing the graph Laplacian from the permuted coordinates, rerunning SpaFlow, and recording the resulting steady state complex concentration 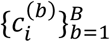 for each spot. We computed a right-tailed rank-based p-value as the fraction of null scores greater than or equal to the original score *c*_*i*_:

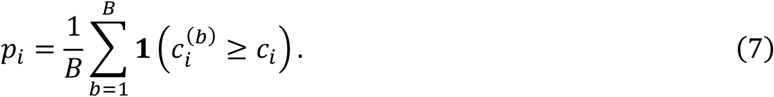

We then applied Benjamini-Hochberg FDR correction within each spot across all tested interactions, and defined hotspots for each LRI as spots with adjusted p-value *q*_*i*_ ≤ *α* (e.g., *α* = 0.05).

### Pathway activity and hotspots definition

Pathway activity was defined by aggregating the activities of all LRIs assigned to a given pathway. Pathway hotspots were defined as the union of hotspots identified from the LRIs within that pathway. According to CellChatDB, all pathways were annotated as secreted signaling pathways. The contribution of each LRI to a given pathway was determined by its number of hotspots.

### Cluster-level CCC via in silico ligand perturbation

To quantify cluster-level CCC from a source cluster *s* to a target cluster *t*, For an LR pair (*l, r*), we performed an in-silico perturbation by setting ligand expression to zero for all spots in *s* while keeping all other spots unchanged and reran SpaFlow to obtain perturbed steady-state complex concentrations. let 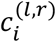 be the original steady-state complex concentration at spot *i* belongs to target cluster *t*, and let 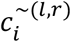 be the corresponding value of cluster *s* after perturbation. We define the CCC strength from *s* to *t* as the average reduction of activity in the target cluster:

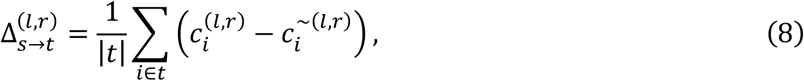

where |*t*| in the number of spots in cluster *t*.

### Evaluation metrics of LRI activity

To benchmark LRI activity inferred by different methods, we evaluated both its spatial organization and its consistency with downstream regulatory programs. We first quantified the spatial autocorrelation of LRI activity using Moran’s I. For each LRI with spot- or cell-level activity scores, Moran’s I was computed on the spatial neighbor graph to assess whether the inferred signal exhibited non-random spatial structure.

We next evaluated downstream regulatory consistency at the cluster level. For each LRI, we averaged its inferred activity across spots/cells within each cluster to obtain a cluster-level LRI activity profile, and then quantified its concordance with downstream regulatory activity across clusters using Spearman correlation. For LRI-TF evaluation, downstream TF activity in each cluster was defined as the mean detection rate of TFs associated with the corresponding LRI, where the detection rate of each TF was calculated as the proportion of spots/cells in that cluster with non-zero expression. For each LRI, we then computed the Spearman correlation between cluster-level LRI activity and cluster-level TF activity across clusters. This detection-based metric was used because TFs are often lowly expressed and sparse in ST data, making detection prevalence more robust than raw expression magnitude. Receptor-TF associations were obtained from scSeqComm^20^ based on the TF_PPR_REACTOME_human resource. For LRI-TG evaluation, downstream TG activity in each cluster was defined as the signed average expression of TGs associated with the corresponding LRI. Specifically, each TG was assigned a regulatory sign according to its TF-target relationship in TRRUST v2^21^: +1 for activation and -1 for repression. For each cluster, TG expression values were multiplied by their assigned signs and then averaged to obtain a signed downstream TG activity score. For each LRI, we computed the Spearman correlation between cluster-level LRI activity and cluster-level TG activity across clusters. Mean expression was used for TGs because they are generally more abundant and less sparse than TFs in ST data, allowing expression magnitude to better capture variation in downstream regulatory output.

### Benchmark tools parameter settings

We conducted a comparative analysis of SpaFlow and other state-of-the-art ST tools for cell-cell communication by evaluating LRI activity using the three metrics described above. For COMMOT, we used the default settings, with ‘min_cell_pct = 0.05’ in the *ct*.*pp*.*filter_lr_database* function and ‘dis_thr = 500’ in the *ct*.*tl*.*spatial_communication* function. LRI activity was defined as the receiver signaling activity inferred by COMMOT. SpatialDM directly infers LRI activity, and we used its default settings to construct the weight matrix. To enable fair comparisons across samples, we did not filter any LRIs in the initial step when calculating global Moran’s R, and we used local Moran’s R, rather than the Fisher’s exact test p-value reported in the original study, as the measure of LRI activity.

### Ablation analysis

For the ablation analysis, we generated four reduced variants of SpaFlow by removing one component at a time from the full model: diffusion, dissociation, production, or degradation. Each variant was applied to the same input data and processed using the same downstream analysis pipeline as the full model. Performance was evaluated using the same three metrics as in the benchmarking analysis and summarized as the mean metric value across all LRIs for each variant. For visualization, metric values were z-score normalized within each sample and displayed as heatmaps to highlight relative performance differences among variants.

### Data analysis for HCC Visium dataset

Six samples from patients with advanced HCC treated with neoadjuvant cabozantinib and nivolumab were reanalyzed in this study. The cohort included three responders and three non-responders (one responder sample was excluded because of low quality). Data preprocessing and spot annotation were performed using the same pipeline described in the original study. Each spot was annotated as “Tumor”, “CAF”, or “Immune”. Tumor boundary spots were defined as non-tumor spots located within three hops of tumor spots. Antigen presentation and T cell scores were calculated using *sc*.*tl*.*score_genes* function in Scanpy^22^ with the corresponding gene sets listed in Supplementary Table 1. The antigen presentation gene set was downloaded from the original study. The T cell gene set included pan-T cell markers such as *CD3D* and *CD3E*, CD8 T cell markers such as *CD8A* and *CD8B*, and CD4 T cell markers such as *CD4* and *IL17A*.

### Significance analysis in mouse heart Visium dataset

The significance of pathway activity between old and young mouse hearts was assessed using a two-sided Wilcoxon rank-sum test followed by Benjamini-Hochberg correction. Significant pathways were defined as those with |log_2_FC| > 0.5 and an adjusted p-value < 0.05. After identifying the eight significant pathways in old mouse hearts, pathway activity was further compared between old and young hearts within each niche using the same statistical testing method. Multiple-testing correction was then applied across niches to determine, for each pathway, which niches showed significant differences. Significance was defined as log_2_FC > 0.5 and an adjusted p value < 0.05, and significant results were marked with ‘*’.

For cell-type proportion analysis, cell-type abundances were obtained from the same deconvolution data used in the original study. Statistical testing was performed as described above by comparing the frequency of each cell type within a given niche with that outside the niche, followed by multiple-testing correction across cell types. Significant differences were defined as log_2_FC > 0.25 and an adjusted p-value < 0.05, and significant results were also marked with ‘*’.

### Correlation between CXCL12-CXCR4 activity and cell-type abundance

Cell-type abundances in human IPF lung Visium data were inferred from matched scRNA-seq data using cell2location^18^. Pearson correlations between CXCL12-CXCR4 activity and cell-type abundance in fibrotic lung spots were calculated separately for fibroblast subtypes and immune subtypes. The top five cell types were selected based on the mean correlation across six samples. Pearson correlation coefficients for each cell type are shown in bar plots, with individual sample values overlaid as dots. The corresponding p-values for each sample are displayed in a heatmap as -log10(p).

### Xenium data analysis

Three samples were included for high-resolution validation: aged control, IPF UL, and IPF LL. Representative Xenium images containing both fibrotic foci and surrounding fibrotic regions were exported from Xenium Explorer to illustrate cell-type colocalization. These illustrative regions were selected based on H&E images and the distribution of CTHRC1+ fibroblasts, with areas showing abundant and spatially clustered CTHRC1+ fibroblasts considered fibrotic foci. CXCL12+ adventitial fibroblasts were defined as adventitial fibroblasts expressing CXCL12 in the Xenium data.

For quantitative analysis, multiple regions were separately and manually annotated as “Foci” (fibrotic foci), “Surrounding” (fibrotic surrounding), or “Other” (normal area) using the same criteria. The “Other” category was annotated only in the aged control sample, whereas the “Foci” and “Surrounding” categories were annotated only in the IPF UL and IPF LL samples. Regions containing fewer than 40 cells were excluded to ensure robust comparison of cell-type fractions. Quantitative comparisons were performed between “Foci” and “Surrounding” regions in IPF UL and IPF LL, and between “Surrounding” regions in IPF samples and the “Other” region in the aged control sample.

## Data availability

Public ST datasets used in this study were obtained from the following sources. The mRCC 10x Visium and CosMx datasets were downloaded from Zenodo under DOI 10.5281/zenodo.16833780. The HCC 10x Visium processed data were obtained from GEO under accession GSE238264; sample HCC3R was excluded because of low cell quality, as indicated by a high proportion of mitochondrial genes. The mouse heart aging 10x Visium dataset was downloaded from Figshare (https://doi.org/10.6084/m9.figshare.28562801.v1), and the matched scRNA-seq dataset was downloaded from Figshare (https://doi.org/10.6084/m9.figshare.28554020.v1). The human IPF Visium, scRNA-seq, and Xenium datasets are not publicly available and can be obtained by contacting the corresponding author.

## Code availability

SpaFlow is an open-source Python package available at https://github.com/Tigerrr07/SpaFlow, code for reproducing the analysis results also available at the same website.

---

**Supplementary Fig. 1.**
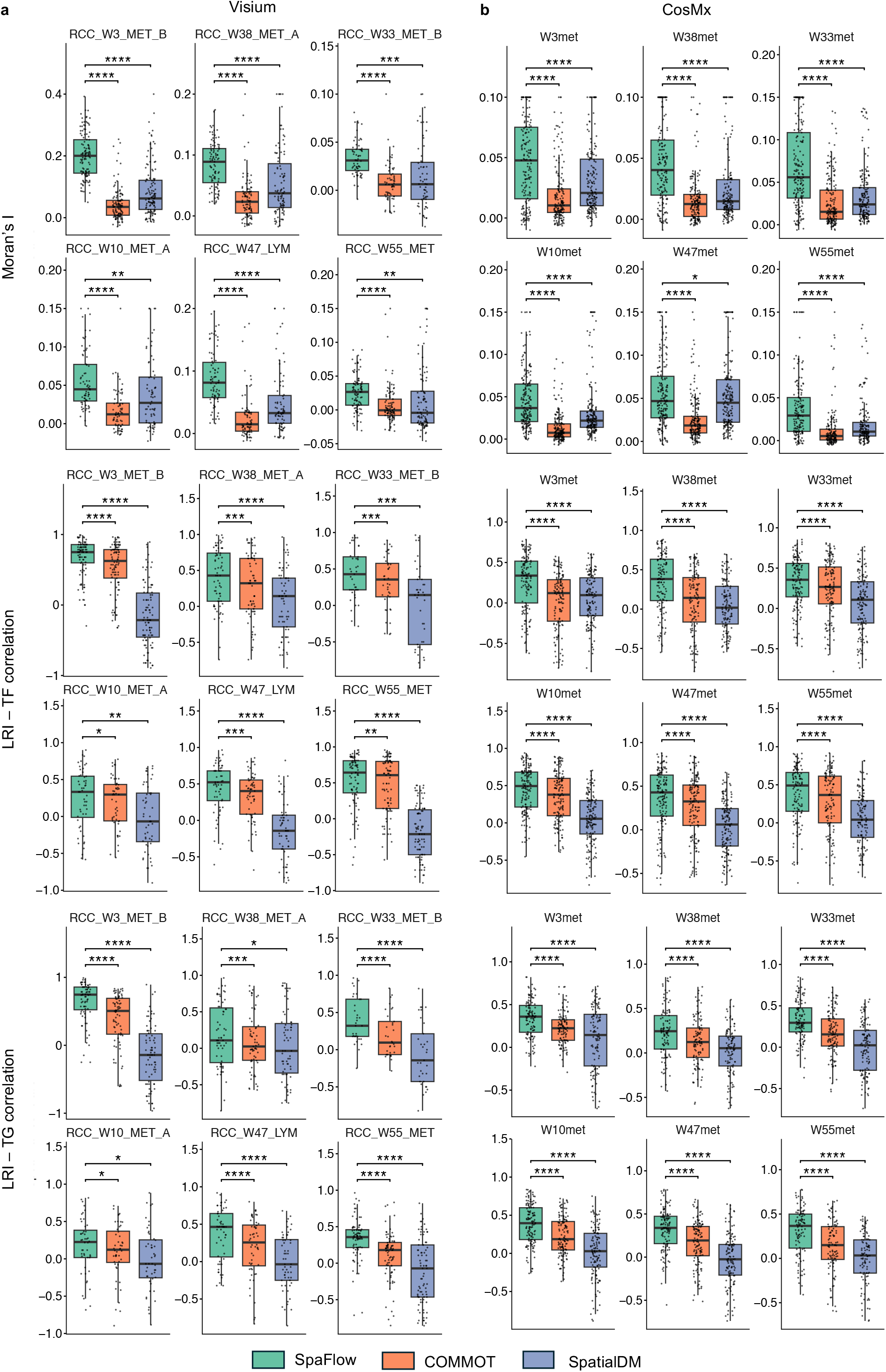
Benchmarking of SpaFlow across all Visium and CosMx samples in the mRCC cohort. **a**, Benchmarking results for all six Visium, comparing SpaFlow, COMMOT and SpatialDM using Moran’s I, and Spearman’s correlations between inferred LRI activity and TF or TG activity. Box plots show the distribution of metric values across LRIs for each sample. Statistical significance was assessed using the Wilcoxon rank-sum test; *P < 0.05, **P < 0.01, ***P < 0.001, ****P < 0.0001. **b**, Benchmarking results across all six CosMx samples using the same three metrics and statistical testing approach as in **a**.

**Supplementary Fig. 2.**
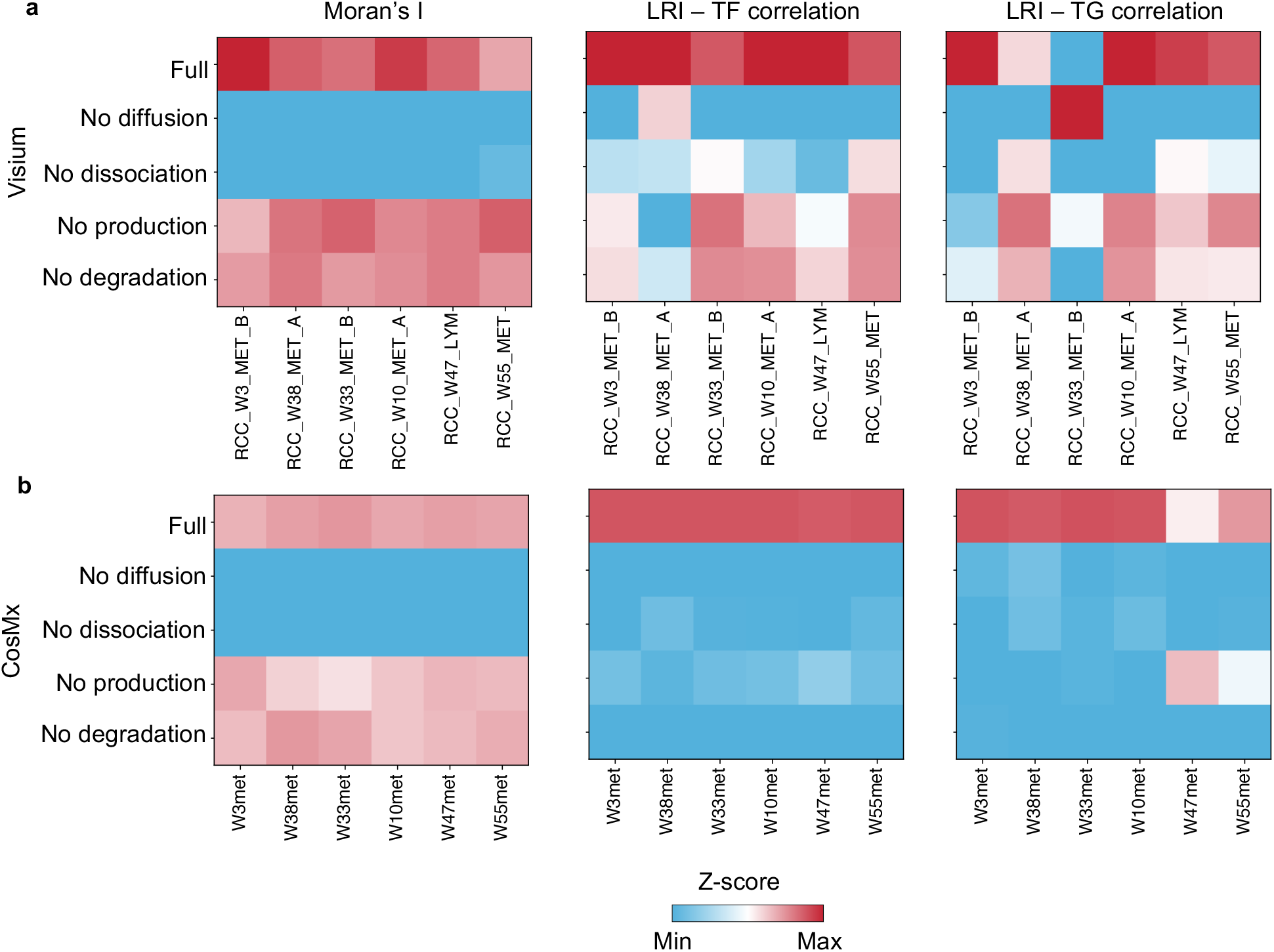
Ablation analysis across all mRCC Visium and CosMx samples. **a**, Heatmaps summarizing model performance across the six mRCC Visium samples. Columns indicate individual samples and rows indicate the full model or ablated models lacking diffusion, dissociation, production, or degradation. Shown are the mean Moran’s I, mean correlation between LRI activity and TF activity, and mean correlation between LRI activity and TG expression, each averaged across all LRIs in each sample. Colors represent Z-scored values within each metric, with red indicating higher and blue indicating lower relative performance. **b**, Same analysis as in **a**, shown for the six mRCC CosMx samples.

**Supplementary Fig. 3.**
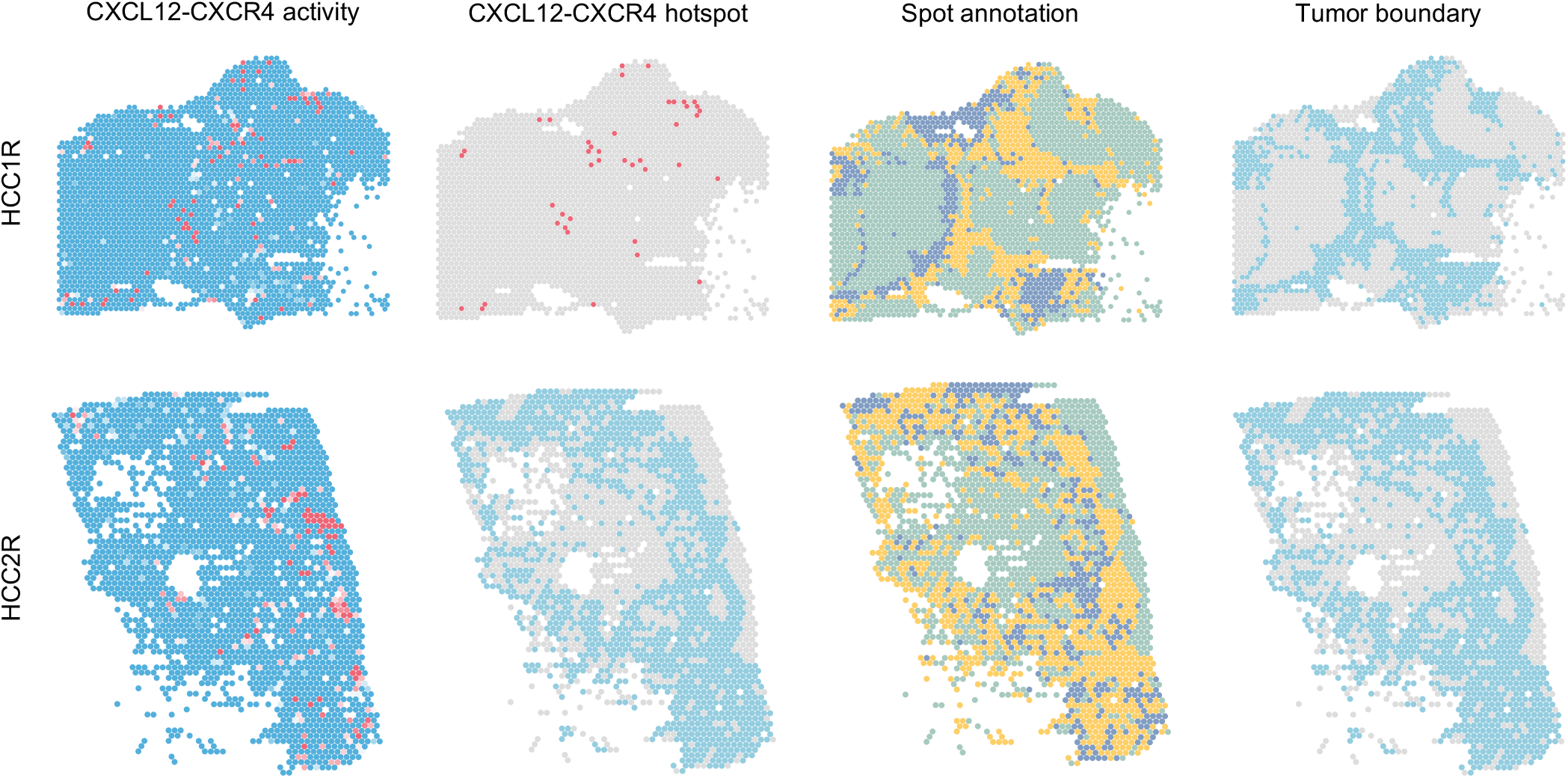
CXCL12-CXCR4 activity and hotspots for responder HCC1R and HCC2R. Spatial maps of HCC1R and HCC2R showing CXCL12-CXCR4 activity, CXCL12-CXCR4 hotspot distribution, spot annotation and tumor-boundary regions.

